# NanoHIVSeq: A Long-Read Bioinformatics Pipeline for High-Throughput Processing of HIV Env Sequences

**DOI:** 10.64898/2026.02.17.706429

**Authors:** Zizhang Sheng, Qin Xiao, Yujie Qiao, Hong Lu, Joseph McWhirter, Manish Sagar, Xueling Wu

**Affiliations:** Aaron Diamond AIDS Research Center, Columbia University Vagelos College of Physicians and Surgeons, New York, NY, USA; Department of Medicine, Division of Infectious Diseases, Boston University Chobanian & Avedisian School of Medicine, Boston, Massachusetts, USA; Department of Virology, Immunology and Microbiology, Boston University Chobanian & Avedisian School of Medicine, Boston, Massachusetts, USA

**Keywords:** *HIV Env*, *nanopore sequencing*, duplex read, viral reservoir, basecalling model

## Abstract

High-throughput sequencing of the HIV-1 envelope (Env) gene from viral quasispecies is essential for epidemiology, virus-antibody coevolution studies, and evaluating therapeutics, but the conventional single-genome amplification (SGA) coupled with Sanger sequencing is labor-intensive and low-throughput. Oxford Nanopore Technology (ONT) offers long-read sequencing advantages, but high error rates (1-7%) poses a challenge in identifying biological variants from sequencing artifacts. Without unique molecular identifiers (UMIs), which lose DNA template and add complexity in library preparation, here we introduce NanoHIVSeq, a UMI-free and reference-free bioinformatics pipeline that processes ONT data from bulk Env PCR amplicons through multistep clustering, consensus polishing, indel correction, denoising, and genotyping to recover functional full-length Env variants. By leveraging advanced ONT duplex sequencing technology, NanoHIVSeq was assessed using plasmid env and bulk HIV reservoir datasets, demonstrating high robustness, recovery rate, reproducibility, and accuracy (>99.9% or >Q30) comparable to UMI approaches. Our findings indicated that NanoHIVSeq allows flexible and simplified ONT library preparation for reproducible and efficient Env sequencing especially for large cohorts.

## Introduction

HIV-1 infects over 1 million people annually which is one of the top concerns of global public health (1). HIV-1 evolves fast in patients and establishes a viral reservoir with high diversity post infection (2, 3). Envelop protein (Env) is a trimeric glycoprotein (160 kD, about 870 amino acids per monomer) that is expressed on the virion surface in charge of cell entry (3). Env is the major antigen that induces host immune response and is thus a target for therapeutic and vaccine development. Circulating HIV-1 has been assigned different groups based on env gene similarity (M, N, O, P). Group M (main) is the most widespread globally and is further divided into 10 major genotypes (A to D, F to H, and J to L) and over 100 circulating recombinant forms (CRFs) (4). Sequencing and genotyping env gene is frequently used for studying HIV epidemiology and virus-antibody coevolution, and for evaluating the efficacy of anti-viral drugs and broadly neutralizing antibodies (3, 5). Conventional approach of sequencing env gene uses env gene PCR amplification from either CD4 T cell genome or plasma, single molecule dilution, and sanger sequencing (6–9). The latter two steps are time-consuming, labor intensive, and costly. The development of third generation sequencing technologies (Nanopore, PacBio, etc.) provides a new opportunity to expedite the sequencing process and reduce the time and cost.

Oxford Nanopore Technologies (ONT) is a third-generation approach that sequences DNA or RNA by measuring changes in electrical current as the molecule passes through a nanoscale protein pore (10). The electrical current signal is converted to nucleotide sequence using dorado or Guppy with machine learning derived basecalling models (10). ONT has been widely applied to sequence viral, plant, and animal genomes due to its advantages including long reads, real-time analysis, portability, scalability, and affordable cost (11). One disadvantage is that ONT reads have high error rates (about 1-7% on average) depending on the basecalling model used (10, 12–14). Many algorithms have been developed to obtain high quality HIV-1 genome sequences and single nucleotide polymorphism (SNP) from ONT reads (reviewed in (15)). Lambrechts et ai. developed a unique molecular identifier (UMI) based approach to sequence HIV-1 provirus from CD4 T cell genome that can identify biological variants with an estimated error rate of less than 0.1% and low indel and PCR recombination rates (12). Another UMI based algorithm, ConSeqUMI was developed to achieve high ONT sequencing accuracy (16). But UMI ONT library preparation requires four or more rounds of PCR and DNA washes. Both previous study and our unpublished data showed that each wash step risk significant DNA loss (17) (e.g., about 10-40% DNA loss per AMPure XP bead purification). This becomes a challenge for sequencing env genes from aviremic samples or donors under antiretroviral therapy (ART), which typically have viral loads of 50 copies or less per mL plasma (18). An additional concern is that sequencing errors in the UMI region reduce the number of usable reads for raw read binning and consensus generation. In addition, large clinical trials recruit hundreds to thousands of donors, the multiple rounds of PCR are still time- and labor-consuming. An optimized protocol with few rounds of PCR, less time consuming, and high accuracy for env gene sequencing is therefore necessary for advancing Env-focused basic and clinical research. Another UMI-free strategy to reduce ONT error rate is to directly align raw reads to generate consensus sequences. Previous studies have explored ONT to generate consensus HIV-1 genomes which showed error rates of 0.1% to 0.6% (13, 14, 19). But these studies obtained a single or limited number of HIV-1 genomes with the goal of identifying drug resistant mutations. Unaddressed concerns of this approach include how to remove PCR and sequencing chimeras, correct indels, and identify all biological variants in a ONT dataset. A UMI-free Env-focused pipeline that can address these concerns will be of great value to the scientific community.

ONT have invented different sequencing and basecalling technologies, but no comprehensive comparison has been performed to determine the optimal settings for obtaining biological variants. For example, ONT R10.4 chip generates simplex and duplex raw reads. Simplex reads are generated by sequencing one strand of the template DNA while duplex reads are generated by sequencing both strands of a template and the signals of both strands are combined to generate a read with higher sequencing quality (quality score (Q) > Q20 or 99% accuracy)(14, 20). To date, for each ONT library from duplex basecalling, the raw reads are a mixture of simplex and duplex reads because duplex sequencing is only successful for a subset of input templates. In addition, ONT uses three machine learning models for basecalling: fast, high accuracy (HAC), and super high accuracy (SUP). The SUP model can obtain the highest basecalling quality score (Q-score) but consumes five-fold more GPU resources than the HAC model. Most previous studies used SUP (10, 12–14), but the optimal combination of basecalling models and ONT read types for biological variant identification is unexplored.

To address above challenges and concerns, we developed a multistep clustering-based bioinformatics pipeline NanoHIVSeq to identify full length functional HIV env variants from UMI-free ONT bulk sequencing dataset. NanoHIVSeq processes ONT datasets prepared from conventional nested-PCR Env DNA and identifies and genotypes functional env sequences. NanoHIVSeq achieves the best performance for determining biological variants using duplex reads from the HAC basecalling model. NanoHIVSeq applies indel correction, denoise, and chimera removal steps to enrich biological variants with a low error rate (<0.05%). Over 90% NanoHIVSeq curated sequences are biological variants or error-free as tested using a library of HIV 32 envs with high diversity (identity <90%) as well as libraries of clinical samples with low diversity (identity >95%). The performance of NanoHIVSeq is highly reproducible and comparable to or better than UMI approaches. Hence, NanoHIVSeq will have broad applications in high-throughput and efficient characterization of HIV-1 env variants.

## MATERIALS AND METHODS

### Reagents

MinION Mk1C sequencing device (MC-114262), and MinKNOW core (6.5.14). Ligation Sequencing Kit V14 (SQK-LSK114) and MinION Flow Cell (FLO-MIN114) were ordered from Oxford Nanopore Technologies. NEBNext Companion Module v2 (E7672S) was purchased from New England Biolabs. Bovine serum albumin (BSA) (50mg/ml) was from Invitrogen (Cat.#. AM2616), and nuclease-free water was from ThermoFisher (Cat. #. J71786-AP), QIAamp viral RNA mini kit (Cat. No. 52904), RNaseOut (Invitrogen, Cat. No. 10777019), SuperScript III (Invitrogen, Cat. No. 18080085), Platinum Taq High Fidelity DNA polymerase (Invitrogen, Cat. No.11304029), ddPCR Super Mix for Probes (No dUTP): BioRad (#1863023), Droplets generated on BioRad DG32 auto generator with associated cartridges (#1864108) and oil for probes (#1863005), Master Mix for Env Droplet PCR: NEB LunaScript Multiplex One-Step RT-PCR kit (E1555L).

### Biological resources

The HIV-1 clade A, B, and C reference rev-env expression plasmids (21–24) were obtained from the NIH HIV Reagent Program as contributed by Julie Overbaugh, Beatrice Hahn, Cynthia Derdeyn, Lynn Morris, and Carolyn Williamson. A total of 32 HIV-1 env plasmids (about 0.5-1 µg each) were pooled and digested by BamHI/XhoI to release the env insert, followed by gel extraction of a near 3kb band for nanopore sequencing.

The plasma samples analyzed in this study were collected from participants in the ADARC Acute and Early HIV-1 Infection Cohort, in which newly infected individuals were recruited and followed longitudinally, with Institutional Review Board (IRB)-approved protocols at the Rockefeller University. All the study cohort participants were infected with clade B HIV-1 (25, 26). At the time of sampling, these subjects had been infected for 3 to 85 months and had not initiated antiretroviral therapy.

### HIV-1 env SGA, bulk PCR, and droplet PCR

The HIV-1 env gene was amplified from patient plasma samples using a single-genome amplification (SGA) method described previously (6–9). Briefly, 140 µl plasma was used to extract viral RNA using the QIAamp viral RNA mini kit (Cat. No. 52904). Reverse transcription (RT) was carried out at 50°C for 60 min, followed by 55°C for an additional 60 min, in a total volume of 100 µl, including 50 µl viral RNA, 20 µl 5x first-strand buffer, 5 µl deoxynucleoside triphosphates (dNTPs) (each at 10 mM), 1.25 µl antisense primer envB3out (27) at 20 µM, 5 µl dithiothreitol (DTT) at 100 mM, 5 µl RNaseOut (Invitrogen, Cat. No. 10777019), and 5 µl SuperScript III (Invitrogen, Cat. No. 18080085). Synthesized cDNA was titrated to single copy, where PCR-positive wells constitute about 30% of the reactions. Nested PCRs were carried out with Platinum Taq High Fidelity DNA polymerase (Invitrogen, Cat. No.11304029. The primers were envB5out and envB3out for the 1st-round PCR and envB5in and envB3in for the 2nd round (28). The cycler parameters were 94°C for 2 min and 35 cycles (45 cycles for the 2^nd^ round) of 94°C for 15 s, 55°C for 30 s, and 68°C for 4 min, followed by 68°C for 10 min. The PCR amplicons were subjected to direct Sanger sequencing, and all sequencing chromatograms were inspected in Sequencher 5.3 (Gene Codes, Ann Arbor, MI) for mixed bases (double peaks), which were evidence of priming from more than one template or of PCR errors and thus excluded. Some 1^st^ round PCRs were pooled for bulk PCRs using the same DNA polymerase system for nanopore sequencing. The Q5 High-Fidelity DNA Polymerase (New England Biolabs) was used for droplet PCR. The pool of first round SGA products was further analyzed using a novel bulk droplet-based PCR methodology that limits recombination (McWhirter et. al., manuscript in preparation). Briefly, the number of templates in the pooled first round SGA amplicons was determined using ddPCR with primers and probes from the Cross-Subtype Intact Proviral Detection Assay (29). The SGA pool was diluted, and around 3,000 templates were distributed in approximately 20,000 droplets along with PCR amplification reagents and 2nd round SGA envelope primers env3Bin and env5Bin. The droplets underwent another 40 amplification cycles before the pooled SGA product was recovered from oil immersions. The recovered PCR product was run on a gel for extraction before nanopore library preparation.

### Nanopore sequencing

We followed the manufacturer’s protocol for Ligation sequencing amplicons V14 (SQK-LSK114). Briefly, the ends of amplicons were repaired and dA-tailed using the NEBNext Companion Module v2. In a 0.2 ml PCR tube, 47 μL amplicon product (about 200 fmol) was mixed with 1 μL DNA control sample of lambda genome, 7 μL NEBNext^®^ FFPE DNA repair buffer v2, 2 μL NEBNext^®^ FFPE DNA repair mix and 3 μL NEBNext^®^ Ultra™ II End Prep enzyme mix. The mixture was incubated at 20 °C for 5 minutes and then at 65 °C for 5 minutes in a thermal cycler. The mixture was then purified by AMPure XP beads. The beads were spined down and washed twice with 200 μL of fresh 80% ethanol. The beads were then air dried for 20–30 seconds and resuspended in 61 μL of nuclease-free water for 2 minutes at room temperature. The eluate was then transferred to a DNA LoBind tube, mixed with 5 μL ligation adapter, 25 μL ligation buffer, and 10 μL Salt-T4 DNA ligase from NEBNext Companion Module v2, and incubated for 10 minutes at room temperature. 40 μL AMPure XP beads was added and incubated for 5 minutes. The beads were then spin down and the supernatant was discarded when clear. The beads were washed twice with 250 μL long fragment buffer and then air dried for 20–30 seconds, resuspended in 15 μL elution buffer, and incubated for 10 minutes. The eluate was then transferred to a new tube. 12 μL DNA library, 37.5 μL sequencing buffer, 25.5 μL library beads was loaded to MinION Mk1C for sequencing following manufacture’s protocol. The 32 plasmid env library and 24 SGA libraries were sequenced using R.10.4.1 chip. The two repeat libraries from donor AD360 month 13 were sequenced using R10.3 chip.

### Bioinformatics analysis

NanoHIVSeq pipeline was used to process nanopore sequencing data. Briefly, NanoHIVSeq calls dorado (v0.9.5) (https://github.com/nanoporetech/dorado?tab=readme-ov-file) to perform basecalling with specified accuracy and model (r1041_e82_400bps_v5.00). By default, Dorado generates simplex reads only. ‘dorado duplex’ command was used to generate duplex reads. In the basecalled bam file, we separated reads to duplex and simplex reads using the dx tag: 0 for simplex while 1 or -1 for duplex or duplex offsprings. NanoPlot (30) v1.42.0 was used to plot sequencing quality of all raw reads and env reads. HMMER v3.4 (31) was used to build an HMM profile for env genes downloaded from CATNAP database. cutadapt v2.7 and seqtk v1.3 were used to filter short reads and separate env, control lambda genome, and undetermined reads using specified primers (env: CAGAAGACAGTGGCA, lambda genome: TGCTTCCAGAGACACCTT). Env reads were converted to forward strand. Usearch (32) v11.0.667 was used to randomly sample up to 1000 raw reads of the lambda genome and Blast (33) v2.10 was used to align raw reads to the lambda reference (genbank accession number: OR974323.1 60-3322) to estimate the proximate error rate of the dataset. Because ONT raw reads contain duplex env reads where the sequencing signals of the two strands failed to combine correctly, nhmmer was used to identify env regions within long raw reads (about 5.2kb). If a raw read contains two env hits with e value <1e-40 and length >2000 nucleotides, both were kept for downstream analyses. Env reads were then clustered using usearch or vsearch (34) v2.30.0 with specified sequence identity cutoff. The clustering includes three steps: the first step is to remove identical reads, reads were then sorted by decreasing sequencing depth or cluster size, and then clustered by prioritizing reads with high sequencing depth as seed at specified identity cutoff (0.995, 0.99, 0.985, 0.98). Clusters containing less than three reads were removed.

Two cycles of racon and 1 cycle of medaka error corrections were performed on each cluster and a consensus sequence was generated per cluster. The consensus sequences were then combined and usearch or vsearch was used to remove duplicates. We observed that about 70% of the unique consensus sequences to have frameshifts which may be caused by sequencing error. Therefore, we developed an alignment-based approach to remove frameshifting indels. Briefly, the unique sequences were aligned using MAFFT v7.52 (35) and a consensus nucleotide was generated for each site. Frameshifting insertions (1, 2, 4, or 5 continuous sites) that were observed in less than 20% of aligned sequences were removed. NanoHIVSeq only corrects insertions with both adjacent sites comprised of over 70% nucleotides but not alignment gaps. Deletions were curated for each sequence separately. Frameshifting deletions of a single nucleotide were replaced by the consensus nucleotide. For deletions of 2, 4, or 5 continuous sites, adjacent nucleotides were removed to ensure the ORF is correct. In frame indels were not corrected. When over three hundred consensus sequences were generated, sequence alignment is time consuming. NanoHIVSeq has an option to first reorder the consensus sequences by sequence similarity and split the sequences to multiple small files (9) with the frameshift correction to be performed on each smaller files without decreasing accuracy. This option can also be applied to datasets containing Envs with high diversity. A set of functional env reads were then identified by removing sequences containing stop codons. Vsearch was used to denoise and remove potential chimeras of the functional envs. A published sliding window method was used to genotyping env variants (36). Briefly, the final set of functional env sequences were fragmented with a 100-nucleotide sliding window. Blastn was used to find the best hit against a reference env dataset presenting all circulating genotypes (459 strains from CATNAP database, 2020 version). The genotype of an env sequence was assigned to the dominant genotype of the fragments. A ‘statistics.txt’ file was generated to summarize the number of reads at each processing step.

Usearch was used to subsample ONT datasets. For pooled SGA ONT library and HIV patient libraries, NanoHIVSeq with the following parameters were used: dorado duplex basecalling using the HAC model; usearch clustering identity cutoff of 0.99; clustering size cutoff of 10.

### Processing published UMI ONT datasets

32 HIV provirus ONT datasets were downloaded from Sequence Read Archive (SRA) database under PRJNA925880. UMI curated consensus sequences were obtained from GenBank with accession numbers MW881651–MW881678, OQ596824–OQ596881, and OR245577– OR246884. These ONT datasets were sequenced with MinION R10.3 flow cells and processed using Guppy with SUP basecalling model (12). Nanoplot reveals an average error rate of 4% for the datasets. Env regions (>1800bp) were extracted from HIV-PULSE curated sequences using HMMER and clustered to remove duplicates using usearch. NanoHIVSeq with sequence identity of 96% was used to cluster raw reads in each of the 32 ONT datasets for consensus read generation and curation. For all ONT runs, curated envs with 10 or more coverage were used for phylogenetic analyses and data size comparison.

The ONT library of 14 plasmid inserts (range of insert size: 404–1884 bp) was downloaded from SRA database (accession number: SRR32256616) (16). This ONT dataset was sequenced with MinION R10.3 flow cells and basecalled using the SUP basecalling model (16). NanoPlot reveals an average error rate of 2.5% for the dataset. ONT raw reads were aligned to references by blastn to remove UMI regions. NanoHIVSeq with sequence identity of 97.5% was used to cluster and curate raw reads.

### Phylogenetic and recombination analysis

Sequences obtained by NanoHIVSeq were aligned using MAFFT and phylogenetic tree was built using IQ-TREE v3 (37–39). Phylogenetic trees were generated by ggtree in R (40). To identify potential chimeras formed by PCR crossover in NanoHIVSeq curated dataset, we used RDP5 (41) to assess potential recombination events and manually examined to identify PCR or sequencing chimeras.

### Statistical analyses

Statistical analyses and plots were generated in R. T-test and paired T-tests were used to compare between NanoHIVSeq settings. Two-sided p-values were calculated. Pearsons’ correlation coefficient was used to determine the correlation between NanoHIVSeq dataset and HIV-PULSE dataset.

### NanoHIVSeq Availability

NanoHIVSeq source code and docker image can be accessed from GitHub: https://github.com/shengzizhang/NanoHIVSeq.

## Results

### The NanoHIVSeq pipeline

To perform nanopore sequencing analysis, we built a bioinformatics pipeline NanoHIVSeq to automate ONT analyses from basecalling to Env genotyping (Figure 1A). Briefly, given an ONT raw read dataset, NanoHIVSeq first removes reads from control DNA and identifies Env region from each raw read and trim off non-Env regions. NanoHIVSeq then clusters Env reads for consensus sequence generation. NanoHIVSeq relies on two primary assumptions to cluster reads for biological variant identification. First, sequencing errors are rare and random events and raw reads with high sequencing depth tend to have low error rates. Using raw reads with high sequencing depth as seeds for clustering and consensus generation is better than random seeds. Thus, NanoHIVSeq applies a two-step clustering method (usearch or vsearch) to group ONT reads by first finding seeds for clustering (reads with high sequencing duplicates) and then clustering all raw reads using a specified sequence identity cutoff. Second, the sequence diversity between two biological variants is increased in ONT reads by an averaged amount of ONT sequencing error rate (Fig. 1C). Thus, clustering ONT raw reads with a sequence identity cutoff close or equal to 1 minus the sequencing error rate can separate reads derived from different biological variants. After sequence clustering, raw reads are polished and one consensus sequence is generated per cluster. Then frameshifting indels in consensus sequences are corrected. An additional denoise step further removes consensus sequences that have low sequencing depth and a high likelihood of containing sequencing errors or PCR chimeras. Then the sequences with correct open reading frame (ORF) are identified as a final dataset of Env variants. Env genotyping is performed on each sequence using a sliding window approach (36).

**Figure. 1.**
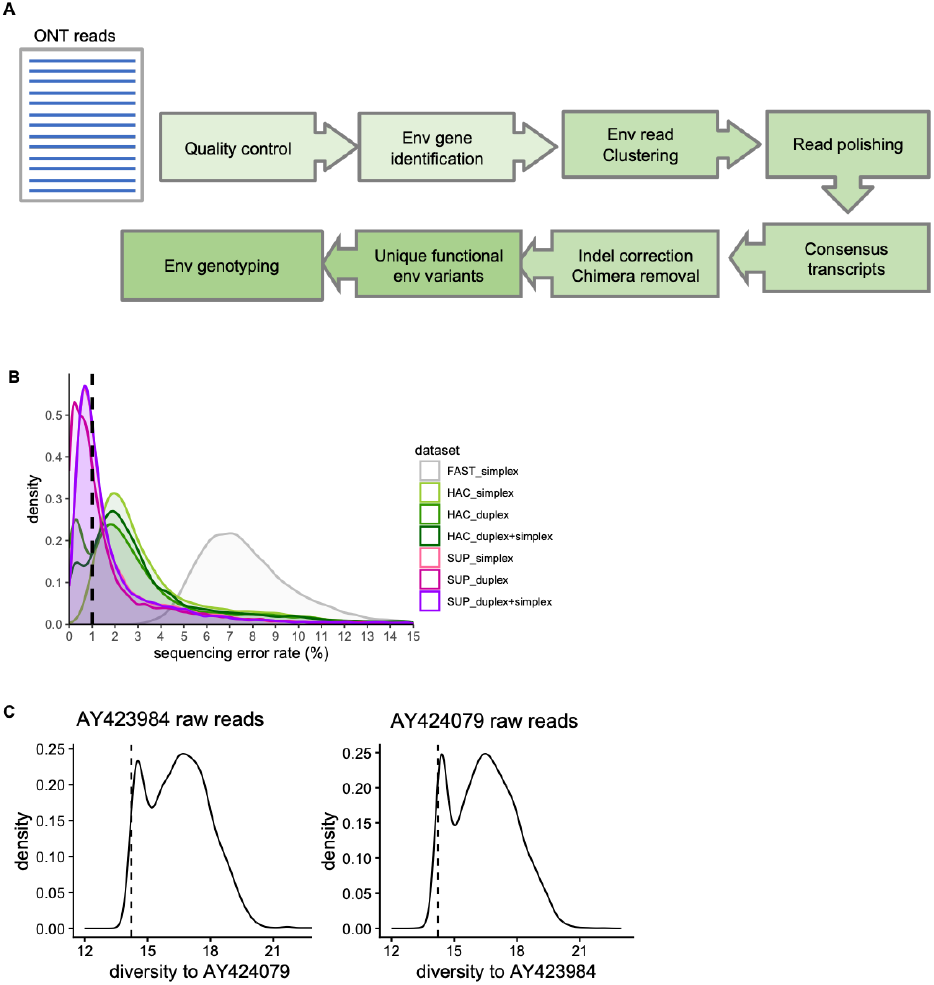
NanoHIVSeq Bioinformatics pipeline and estimated sequencing errors. (A) NanoHIVSeq data processing steps. (B) Comparison of sequencing error rates of duplex and simplex reads from fast, HAC, and SUP basecalling modes. ‘duplex+simplex’ represents all reads generated from duplex basecalling while ‘simplex’ represents simplex basecalling. Note: a few raw reads with over 15% errors were not shown. (C) Diversity of AY423984 ONT raw reads to AY424079 (left) and diversity of AY424079 ONT raw reads to AY423984. HAC duplex reads were used for the plots. The diversity between AY424079 and AY424079 was shown as dashed line.

### Optimization and performance assessment of NanoHIVSeq

We hypothesize that sequencing error rate, ONT read types, basecalling model, and clustering methods contribute to the performance of NanoHIVSeq. To optimize NanoHIVSeq, we sequenced a ONT library of published Env genes (Fig. S1A). Briefly, 32 Envs with high diversity (mean identity: 82%) were pooled and released from plasmids for direct ONT sequencing (Figure S1B). ONT reads were basecalled using three basecalling models (fast, HAC and SUP) and two basecalling methods (simplex and duplex), yielding the following datasets: fast simplex dataset, HAC simplex dataset, SUP simplex dataset, HAC duplex dataset, and SUP duplex dataset (see Methods). Note, as described in the Introduction, the duplex dataset is comprised of duplex and simplex reads. For each duplex dataset, we further compared duplex reads only (referred to HAC duplex and SUP duplex in Fig. 1B) and the mixture (HAC ‘duplex+simplex’ and SUP ‘duplex+simplex’ in Fig. 1B). For all analyses below, HAC duplex and SUP duplex refer to duplex reads only dataset. In total, we compared sequencing quality of seven datasets (Fig. 1B). Overall, we obtained about 500,000 to 750,000 ONT reads for each basecalling model (Figure S1C). The SUP model generated about 5% more reads than the HAC model (Figure S1C). Each dataset contains about 30-60 percent Env reads (Figure S1C). We aligned the Env reads to the references and calculated the sequencing errors per dataset. SUP duplex reads showed the lowest error rate (<0.5%) followed by SUP ‘duplex+simplex’ and SUP simplex dataset. HAC duplex reads showed two error rate peaks: less than 0.5% and about 2% (Figure 1B) while HAC ‘duplex+simplex’ and simplex datasets showed higher 2% error peaks. FAST model-derived Env reads contained an average of 7% errors which is too high to be used for downstream analyses. The error rate distributions of Env gene are similar to those of control DNA (Figure S2A) except that the control DNA have a higher proportion of ONT reads with fewer errors (Figure S2B). Next, we applied NanoHIVSeq to six datasets above except fast simplex to identify the reference Env sequences and compared the performance of combination of different settings including clustering method (usearch vs vsearch), basecalling models, read types, and clustering cutoffs (sequence identity: 0.980, 0.985, 0.990, and 0.995). Because a subset of reference Envs were sequenced with low depth (Fig. S1D), not all 32 Envs can be identified from the six datasets. 30 Envs have over ten raw reads. By using the final curated Env sequences from each dataset, the performance of different settings was first evaluated by three indices: ratio of recovered Env strains (R_rs_ =number of recovered strain/total number of reference Envs), ratio of biological variants or variants identical to reference (R_bv_ = number of biological variants /number of total variants), and mean number of variants per reference (M_vr_ =number of total variants/number of recovered references). The optimal pipeline should have a high R_rs_ and R_bv_ and a low M_vr_ . Overall, for all six datasets, usearch and vsearch showed comparable R_rs_ (Fig. 2A) but usearch had a significantly higher R_bv_ (Fig. 2B), indicating that usearch clustering is better at enriching biological variants. We therefore used reads clustered by usearch for consensus generation. Analysis of the open reading frame (ORF) of the consensus sequences showed an overall 30% sequences to have correct ORF (Fig. 2C). Majority of the rest sequences tend to have frameshifting indels or in frame stop codons. We developed an alignment-based approach to screen and correct frameshifting indels which increased sequences with correct ORFs to 80% (Fig. 2C). After indel correction, the vsearch sequence denoising and chimera removal module was applied to examine whether a sequence with low sequencing depth are erroneous copies of sequences with high sequencing depth. The module improved both M_vr_ and R_bv_ by over three to four-fold without losing biological variants (Fig. 2D and S2C to S2D). The comparison of final curated Envs revealed that optimal combination of the three indices was obtained from HAC duplex with clustering cutoff 0.990, HAC ‘duplex+simplex’ with clustering cuoff of 0.995, and HAC duplex with clustering cutoff of 0.995 (Fig. 2E). For these three datasets, we further compared their performance at three minimum clustering sizes (3, 10, and 15 ONT reads). Curated Envs from the HAC duplex dataset with a 0.990 clustering cutoff showed a R_bv_ comparable to the other two datasets but better R_rs_ and M_vr_ at minimum clustering coverages of 10 and 15 reads (Fig. 2F). Thus, these results indicated that curated Envs from the HAC duplex dataset with a 0.990 clustering cutoff and minimum clustering sizes of 10 and 15 reads were optimal for the three indices.

**Figure. 2.**
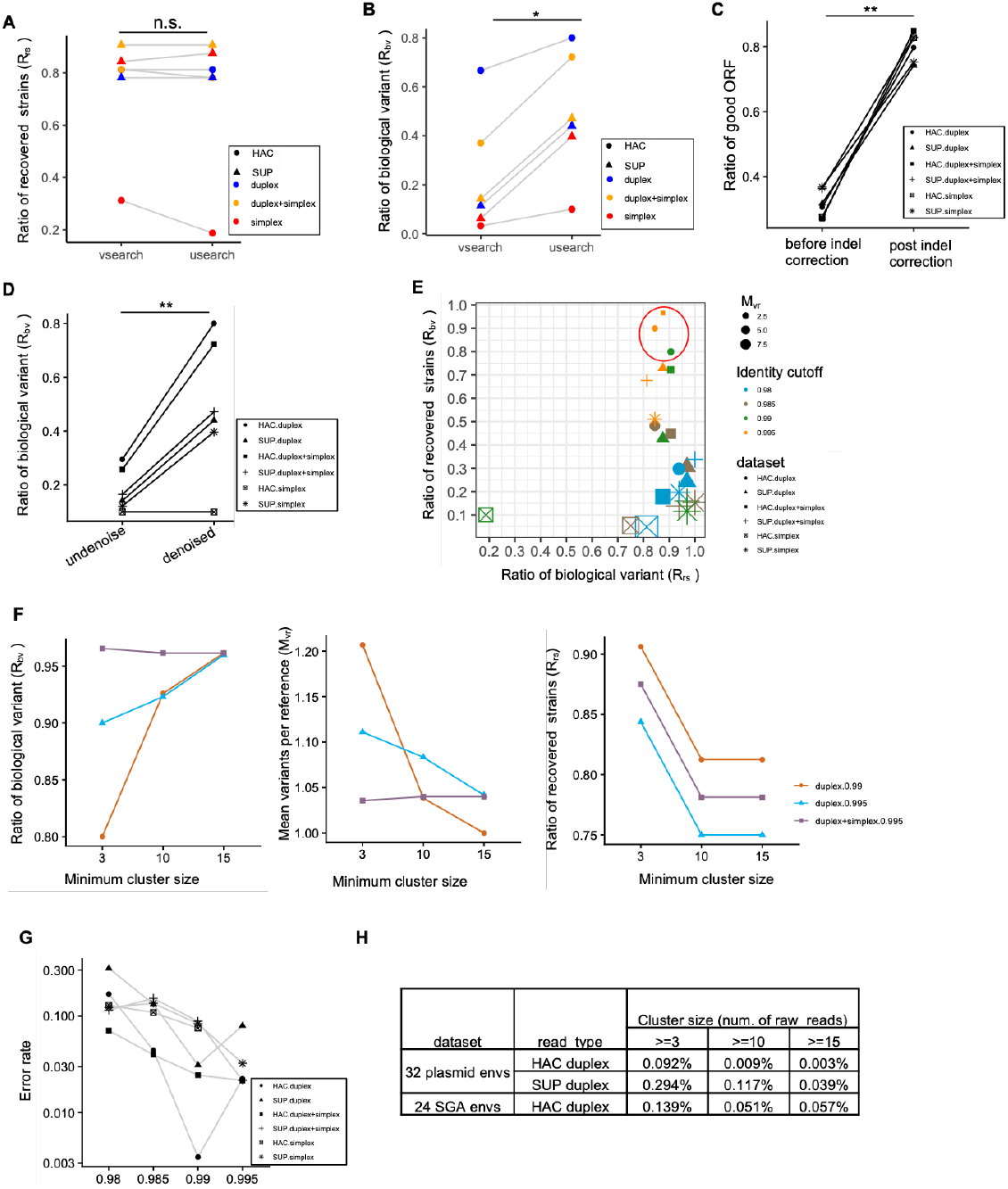
Comparison of clustering software and basecalling models on ONT accuracy. (A) The total reference envs found by usearch and vsearch clustering are comparable using ONT reads generated by different basecalling models. . Paired T-test : p-value = 0.54. (B) Usearch clustering generates less consensus sequences using ONT reads generated by different basecalling models, resulting in a higher ratio of sequences identical to reference (ratio of biological variants) than vsearch. Paired T-test : p-value = 0.03. (C) The NanoHIVSeq indel correction module significantly improved the rate of sequences with correct open reading frame. Paired T-test: p-value <0.0001. Usearch clustering with identity cutoff of 0.99 was used to generate consensus sequences. (D) Denoise and chimera filter substantially reduced the number of variants per reference strain. Usearch clustering with identity cutoff of 0.99 was used to generate consensus sequences. Paired T-test : p-value = 0.008. (E) Comparison of different ONT basecalled dataset and sequence identity clustering cutoff on R_rs_, R_bv_, and M_vr_ . (F) Consensus sequences generated from larger clusters have improved R_bv_ and M_vr_ but reduced R_rs_ . (G) Sequencing error rate decreased when using higher sequence identity cutoff for consensus sequence generation. (H) Sequencing error is reduced when sequencing depth increase. For both SUP and HAC reads, sequencing error is reduced substantially when sequencing depth increase to over 10 reads.

By applying the optimal settings determined above, we calculated the error rate of the curated Envs and investigated the impacts of cluster size and clustering cutoff on sequencing error rate. For each of the six ONT datasets, we calculated sequencing error for each curated Env sequence. Overall, lower error rates were observed for Envs curated from higher clustering identity cutoff and higher minimum cluster size (10 and 15 ONT reads) (Fig. 2G, S2E, and S2F). The HAC duplex and HAC ‘duplex+simplex’ datasets have lower error rates than Envs curated from other datasets (Fig. 2G, S2E, S2F, and S2G). Curated Envs from the HAC duplex dataset have the lowest error rate at clustering cutoff 0.99 and minimum cluster sizes of 10 and 15 reads (error rates: 0.009% and 0.003% respectively, Fig. 2H). The quality of the curated Envs are greater than Q40, which are comparable to those of UMI approach when sequences were curated at similar bin sizes (12). These results indicated that applying NanoHIVSeq to HAC duplex dataset obtained the optimal results measured by the three indices and error rate.

To further understand the robustness of NanoHIVSeq’s performance, subsampling from 10,000 to 300,000 reads of the HAC duplex dataset was performed. For minimum clustering sizes of 10 and 15 reads, M_vr_ and R_bv_ converged to around 1.05 and 0.96 respectively when subsampling 100,000 or more reads; R_rs_ reached around 0.80 (Fig. S3A). All three indices tend to converge slower for clustering size cutoff of 3. This suggested that both size cutoffs 10 and 15 are more robust to input read numbers. The Env gene error rates of clustering size cutoffs of 10 and 15 reads were consistent with those obtained from the full HAC dataset (0.011%+0.008% and 0.0089% +0.0095%) (Fig. 2H, S3B and S3C). In addition, previous studies used SUP reads for consensus sequence generation, the subsampling analysis showed that the error rates of consensus sequences generated from SUP duplex reads were higher or comparable to that of the HAC duplex reads (Fig. S3B and S3C). Curation of the SUP duplex dataset showed comparable M_vr_ and R_rs_ to HAC duplex dataset but a significantly worse R_bv_ (Fig. S3D), indicating that SUP reads may not benefit NanoHIVSeq’s performance.

In summary, our results revealed contributions of diverse settings and their optimal combination for NanoHIVSeq to obtain high confident biological variants. These optimal settings include HAC duplex reads for input, usearch for clustering with 0.99 as clustering identity cutoff, vsearch for denoise and chimera removal, and clustering size of 10 or more reads. With the optimal parameters, over 90% of NanoHIVSeq curated Envs are identical to the references while the rests have a low error rate. 26 of the 30 Envs sequenced with more than 10 raw reads were recovered. RDP5 did not find sequencing chimeras from the curated Env sequences.

### Performance assessed using HIV-1 reservoir libraries

To assess the performance of NanoHIVSeq on HIV-1 variants, we performed ONT sequencing on two libraries. In the first library, we pooled 24 Envs derived from single genome amplification (SGAs) of two HIV-1 patients (AD360 and AD415). Within each donor, SGAs showed greater than 95% Env sequence identity (Fig. S4A) which reflects intra-donor Env diversity. The first round PCRs of these SGAs were pooled and PCR amplified and then subjected to ONT, repeated for three times (Fig. S4B). The three repeats showed high similarity in sequencing coverage for the references (Fig. S4C), with a total of 13 references detected (Fig. S4C). Each ONT repeat contains eight or nine references identifiable by NanoHIVSeq (Fig. S4C). The results showed high overlap of the curated Envs from the three ONT repeats (Fig. 3A). M_vr_, R_bv_, R_rs_ and error rate were comparable to those of the 32 plasmid Env dataset (Fig. 2H and 3B). Phylogenetic tree revealed high similarity to the Sanger sequenced SGAs (Fig. 3C).

**Figure. 3.**
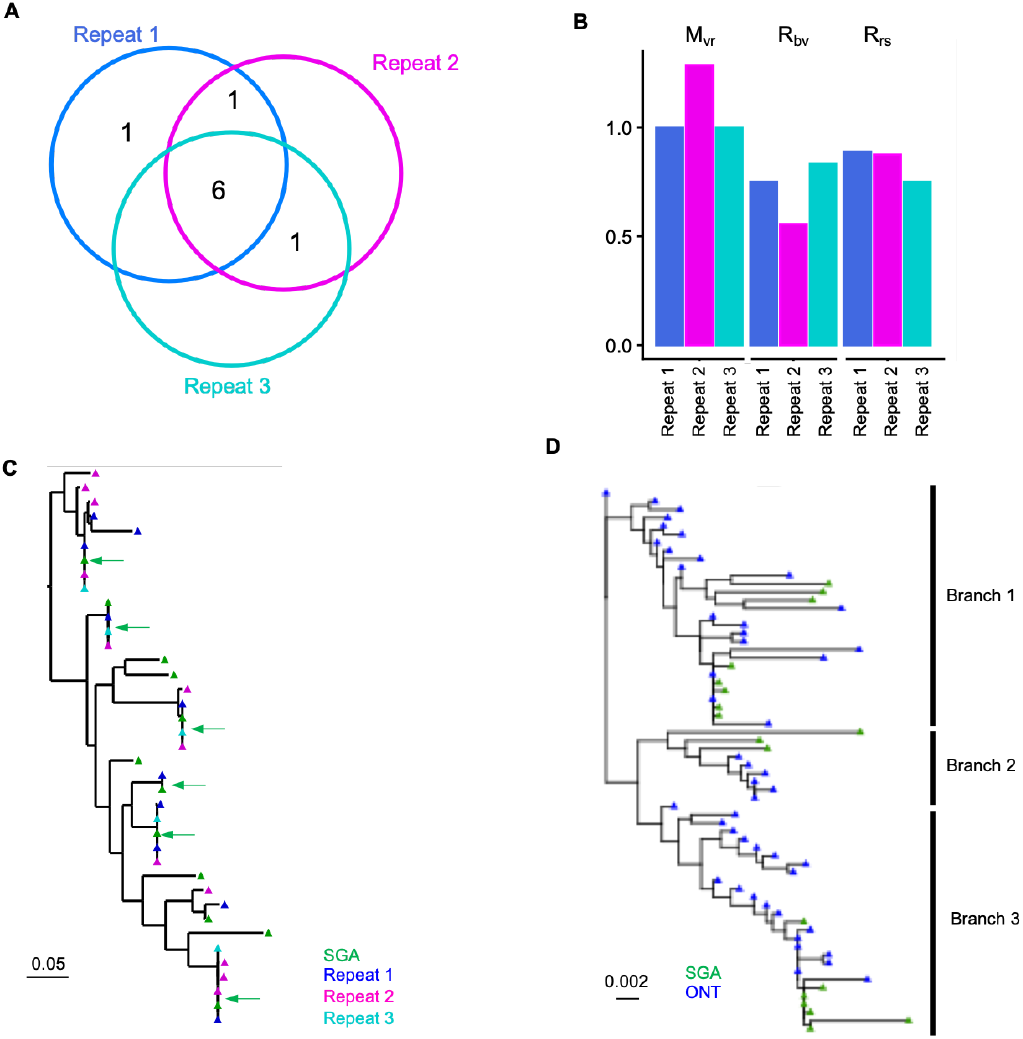
Assessment of reproducibility and performance of NanoHIVSeq using SGA pool and HIV-1 reservoir ONT dataset. (A) Shared env sequences between three ONT runs of 24 SGAs with high sequence similarity. (B) R_rs_, R_bv_, and M_vr_ indices of the three ONT repeats of the SGA library sequencing. (C) Phylogenetic tree of AD360 env sequences obtained from three ONT repeats showed high reproducibility. Green arrows showed curated ONT sequences identical to SGA sequences. (D) Comparison of env sequences obtained from ONT reads to SGA-derived env sequences reveals high reproducibility between ONT repeats and SGA sequences. ONT curated reads revealed a detailed pathway of env evolution.

The second library was from donor AD360, where four 1^st^ round Env PCRs from multiple copies of viral genome were pooled for bulk PCR and ONT sequencing. NanoHIVSeq revealed 48 curated functional Envs. Env cDNAs were also diluted for single genome sequencing and 18 variants were obtained. The phylogenetic tree revealed that Env gene evolved three branches (Fig. 3D). Envs from ONT and SGAs clustered together across the three branches. ONT Envs revealed more detailed pathways of Env diversification between the three branches. In summary, the results from both libraries demonstrated that NanoHIVSeq coupled with ONT can sequence Env variants with high accuracy and reproducibility and recapitulate Env evolutionary pathway in HIV-1 patients.

### Comparison of NanoHIVSeq to UMI approaches

We compared the performance of NanoHIVSeq to two UMI approaches: HIV-PULSE and ConSeqUMI. In Lambrechts et al study (12), a total of 1308 HIV-1 proviruses were obtained from eighteen donors using 32 ONT datasets with HIV-PULSE. In these provirus sequences, a total of 699 Envs greater than 1800bp were obtained with 633 being unique. NanoHIVSeq was applied to the 32 ONT datasets to find Env regions greater than 1800bp and generate curated Envs. 6526 and 1703 curated Envs were obtained for minimum clustering sizes of 3 and 10 respectively. NanoHIVSeq curated dataset contained more Envs than those found in the study for three primary reasons. First, NanoHIVSeq clustered all HIV-1 reads in each ONT run for curation while HIV-PULSE used about 67% HIV-1 reads with confident UMI binning. Second, HIV-PULSE used a minimum bin size of 15 reads which was higher than NanoHIVSeq. Third, Env amplicons from each donor were sequenced in over five ONT runs. The Lambrechts et al study further clustered Envs with a 99.5% sequence identity from all ONT runs per donor and curated Envs. To demonstrate the reproducibility of NanoHIVSeq, we pooled curated Envs from all ONT runs without further clustering. Overall, 92%, 83% and 52% of the 633 HIV-PULSE curated Envs were observed in the NanoHIVSeq curated dataset with >99%, >99.9%, and 100% sequence identities respectively (Fig. 4A). The total number of curated Envs per donor was significantly correlated between HIV-PULSE and NanoHIVSeq curated datasets (Pearsons’ r: 0.60, p=0.007) (Fig. 4B). Phylogenetic trees were constructed for Envs in each donor and revealed high reproducibility between HIV-PULSE and NanoHIVSeq curated datasets as well as between NanoHIVSeq curated Envs from different ONT runs (Fig. 4C, S5A-S5D). RDP5 detected 4 PCR or sequencing recombination from the 6526 Envs, indicating a recombination rate of about 0.06%. All 4 chimeras were curated from clusters with sizes less than 10 which can be effectively excluded by setting the clustering size cutoff to 10.

**Figure. 4.**
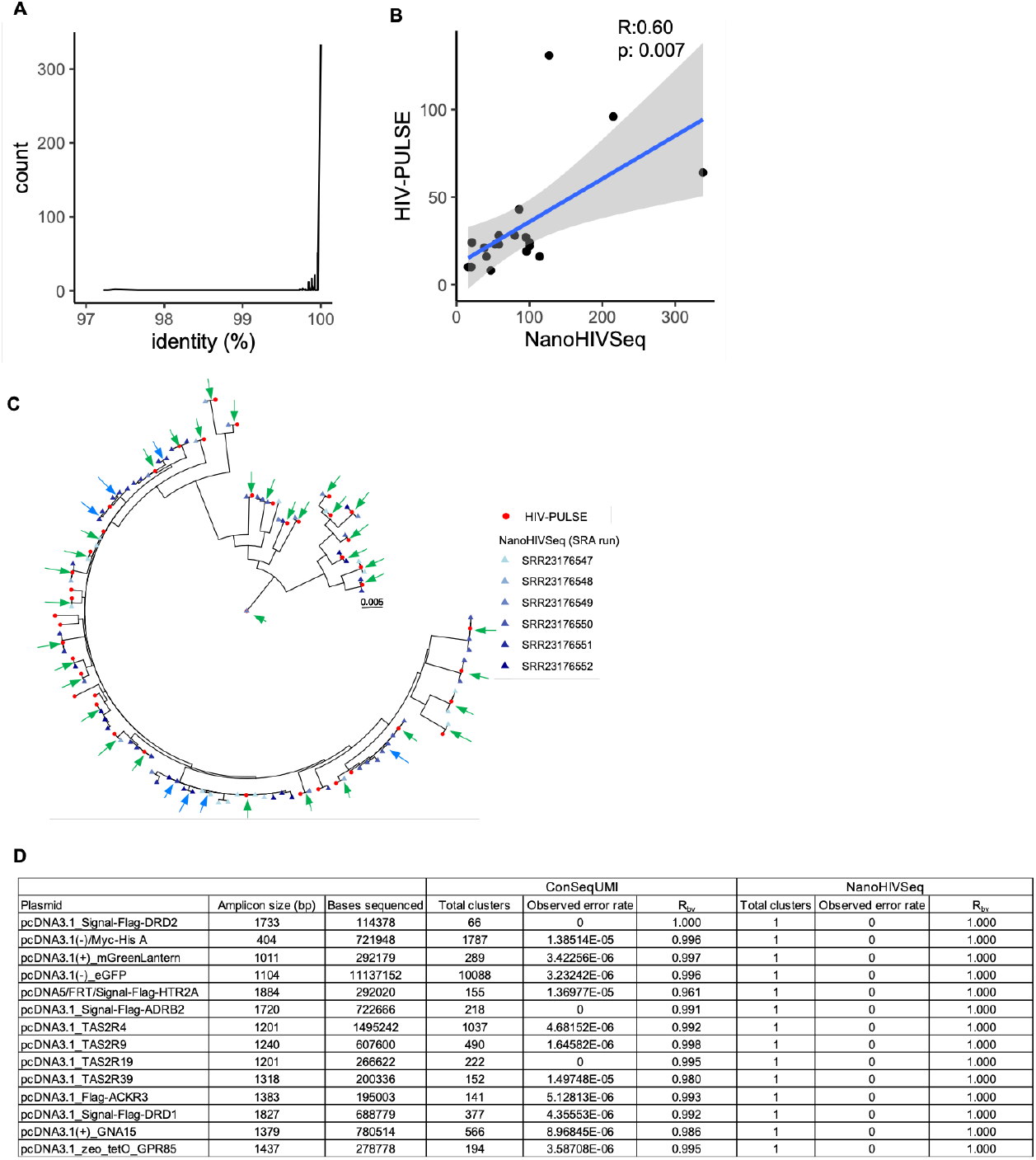
Comparison of NanoHIVSeq to UMI approaches HIV-PULSE and ConSeqUMI. (A) Sequence identity distribution of HIV-PULSE generated env sequences to the most similar sequences in NanoHIVSeq curated env sequences. (B) Correlation of per donor data size between NanoHIVSeq and HIV-PUSLE curated env datasets. Person’s r and p values were shown. (C) Phylogenetic tree showed high reproducibility of NanoHIVSeq and HIV-PULSE generated env sequences from donor MRC008. Green and blue arrows showed identical sequences between NanoHIVSeq and HIV-PUSLE and bweteen NanoHIVSeq ONT runs respectively. (D) Comparison of cDNA generated by NanoHIVSeq and ConSeqUMI showed that NanoHIVSeq is effective at finding biological sequences with no sequencing error. ONT dataset information was obtained from the Zahm et al study.

To assess the performance of NanoHIVSeq on ONT libraries other than HIV-1 Env, we applied NanoHIVSeq to a ONT library comprised of 14 plasmid inserts sequenced with high depth by Zahm et al (16). Overall, NanoHIVSeq generated one curated sequence per plasmid insert (Fig. 4D). These curated sequences were identical to the references. In comparison, the ConSeqUMI approach generated more clusters than NanoHIVSeq. Both approaches recovered the 14 inserts (R_rs_ =1). The R_bv_ per cDNA insert ranged from 0.96 to 1.0 for ConSeqUMI curated dataset while the R_bv_ of NanoHIVSeq curated inserts are 1.0. In total, ConSeqUMI curated references consisted of 88 unique variants corresponding to an M_vr_ of 6.3 compared to Mvr of 1.0 for NanoHIVSeq curated inserts. This highlighted that NanoHIVSeq can be used to process ONT libraries other than Env gene with high accuracy.

In summary, our results showed that NanoHIVSeq generated high quality biological variants from ONT sequencing with high reproducibility and accuracy. The performance of NanoHIVSeq is at least comparable to the two state-of-the-art UMI approaches HIV-PULSE and ConSeqUMI.

## Discussion

High-throughput sequencing of HIV Env variants is needed for evaluating antibody-based therapeutic effectiveness in clinical trials and epidemiological surveillance. In this study, we developed a UMI-free and reference-free pipeline, NanoHIVSeq, to process ONT datasets for identification of biological variants with high accuracy and reproducibility. We optimized and assessed the performance of NanoHIVSeq using both heterogeneous and homogeneous HIV Env libraries sequenced by the most advanced ONT duplex sequencing technology (R.10.4 flow cell). The performance of NanoHIVSeq was further assessed using ONT datasets with higher error rates (provirus and cDNA insert libraries, R.10.3 flow cell). NanoHIVSeq showed a performance comparable to or better than the UMI approaches as well as published UMI-free approaches (12, 15, 16, 19, 42).

Obtaining error free or biological sequences from ONT dataset is the goal for ONT algorithm development. In this study, we used three indices to assess the performance of NanoHIVSeq: R_rs_, R_bv_, and M_vr_ . R_bv_ measures the percent of biological variants in the curated sequences. Our analyses revealed that NanoHIVSeq achieves a R_bv_ over 0.9 to 1.0. R_rs_ measures the total number of references recovered irrespective of sequencing error. R_rs_ is sequencing depth dependent. We observe that a sequencing depth of 10 raw reads or more is needed for high confidence. M_vr_ reveals the capability of an algorithm to identify biological references from erroneous references. We showed that the indel correction module and vsearch denoise and chimeral removal is critical to correcting and excluding erroneous sequences. Although PCR and sequencing induced recombination is a key issue for Env sequencing, we demonstrated that NanoHIVSeq curated sequences have a very low rate of sequence chimeras. To further reduce sequencing chimeras, users can prepare ONT libraries with protocols using droplet PCR and emulsion PCR (29). With less PCR steps and DNA washes than UMI approaches, NanoHIVSeq coupled with ONT may enhance the success rate of sequencing aviremic samples, which is now under investigation.

To date, the current ONT flow cell 10.4 produces duplex reads that have higher sequencing quality than simplex reads, which were the only options in the earlier generations of flow cells. But few studies have assessed the performance of duplex and simplex reads as well as their combination with HAC and SUP basecalling models in biological variant curation. Our results showed that the optimal results can be obtained by HAC duplex reads. A sequencing depth of 10 HAC duplex raw reads are enough to curate references with 0.05% or less errors, which is about 1 nucleotide error per Env. Majority of curated sequences with larger cluster sizes are error free (Fig. S2G). Because the HAC model basecalling is about 5 times faster than the SUP model, this result revealed a more efficient way of ONT data processing. Future investigation is needed to examine if HAC duplex reads can be used to improve UMI approaches.

NanoHIVSeq has the following limitations. First, the sequence clustering identity cutoff is ONT sequencing error rate dependent. We applied cutoffs of 0.99, 0.96, and 0.975 for the ONT duplex datasets, provirus datasets, and plasmid insert dataset respectively. Our analyses revealed that lowering the identity cutoff will not affect the recovered biological variants (Fig. 2E) but may introduce more curated sequences with low cluster sizes. In such cases, removal of curated sequences with low cluster sizes is needed to filter out low quality sequences. A clustering identity cutoff higher than the optimal may generate few sequence clusters. Nonetheless, we observed that the optimal cutoff is close to 1.0 - the error rate of the raw dataset which can be calculated by Nanoplot. Second, NanoHIVSeq curated datasets may not be used to study viral clonal expansion, an issue also presented in UMI approach (12). This is because NanoHIVSeq has multiple sequence duplicate removal steps to minimize sequencing errors in the final dataset. An alternative approach is to sequence viral DNA in multiple ONT runs and use identical or highly similar sequences among ONT runs as evidence of clonal expansion.

In summary, NanoHIVSeq can process both ONT datasets generated from different amplicon protocols. NanoHIVSeq improves ONT library sequencing efficiency and is an ideal alternative approach for sequencing large cohorts.

## Data Availability

ONT raw reads were deposited to SRA database https://www.ncbi.nlm.nih.gov/sra/ with project ID: PRJNA1419998. SGA and NanoHIVSeq curated sequences were deposited to GenBank https://www.ncbi.nlm.nih.gov/genbank/. NanoHIVSeq source code is available at GitHub: https://github.com/shengzizhang/NanoHIVSeq. Both NanoHIVSeq curated sequences and source codes are available at Figshare with DOI: 10.6084/m9.figshare.31347640.

## Supplementary Data statement

Supplementary Data are available at NAR Online.

## Author contributions

Z.S. and X.W. designed the research. Z.S. and Y.Q. wrote the methods and analyzed the data. Q.X. performed ONT sequencing. J.M., H.L., X.W., M.S. prepared the ONT libraries. Z.S., X.W., and Q.X. wrote the paper, and all authors reviewed, commented on, and approved the manuscript.

