## supplemental_figs for "NanoHIVSeq: A Long-Read Bioinformatics Pipeline for High-Throughput Processing of HIV Env Sequences"

**A**

| HIV Strain | Accession Number |
|---|---|
| 246_F3_C10_2 | HM215279 |
| 398_F1_F6_20 | HM215312 |
| 6535.3 | AY835438 |
| BJOX002000.03.2 | HM215364 |
| CAAN5342A2 | AY835452 |
| CAP45.2.00.G3 | DQ435682 |
| CE1176_A3 | FJ444437 |
| CE703010217_B6 | FJ443575 |
| CH119.10 | EF117261 |
| CNE55 | HM215418 |
| CNE8 | HM215427 |
| Du123.6 | DQ411850 |
| Du156.12 | DQ411852 |
| Du172.17 | DQ411853 |
| HIV_25710-2.43 | EF117271 |
| PV0.4 | AY835444 |
| QH0692.42 | AY835439 |
| RE-104541.67 | AY835449 |
| RHPA4259.7 | AY835447 |
| SS1196.1 | AY835442 |
| THR04156.18 | AY835448 |
| TRJ04551.58 | AY835450 |
| TRO.11 | AY835445 |
| WIT04160.33 | AY835451 |
| X1632_S2_B10 | FJ817370 |
| ZM109F.PB4 | AY424138 |
| ZM135M.PLIO | AY424079 |
| ZM197M.PB7 | DQ388515 |
| ZM214M.PL15 | DQ388516 |
| ZM233M.PB6 | DQ388517 |
| ZM249M.PL1 | DQ388514 |
| ZM53M.PB12* | AY423984* |

**B**

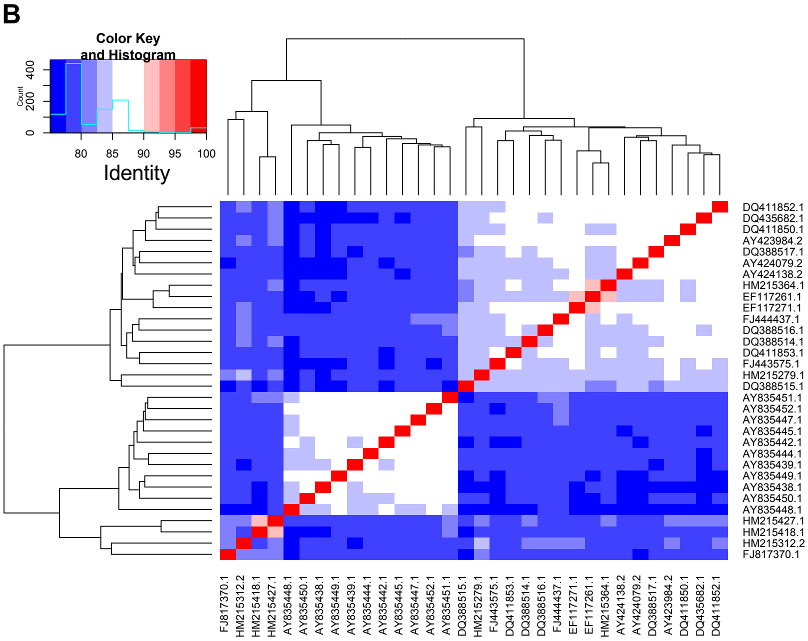


**C**

| Read type | Calling model | Rawreads | Duplex_reads | Rawread_spikein | ENV_raw | ENV_unique |
|---|---|---|---|---|---|---|
| Simplex | FAST | 634474 | NA | 13132 | 355564 | 355564 |
| | HAC | 506872 | NA | 17122 | 434374 | 433643 |
| | SUP | 526353 | NA | 18405 | 451143 | 293051 |
| Duplex | HAC | 711915 | 332353 | 5803 | 300857 | 233765 |
| | SUP | 748999 | 388631 | 6074 | 346102 | 181357 |

**D**

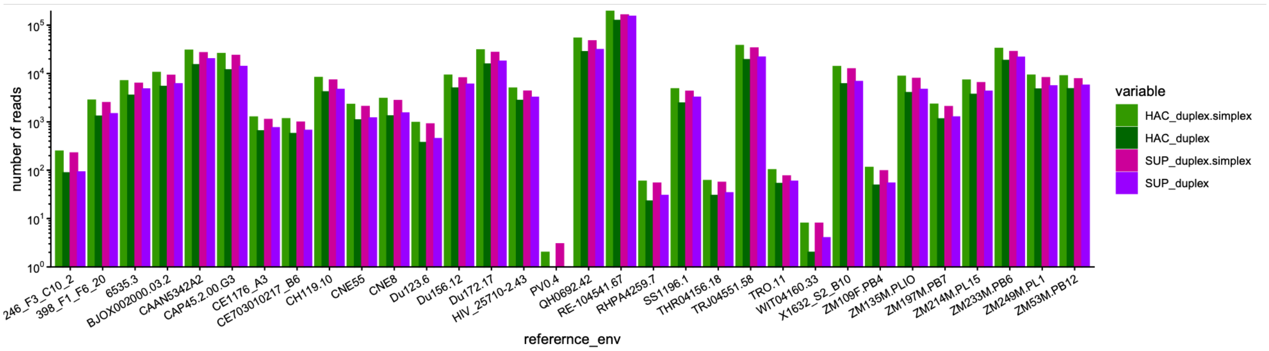


**Figure.S1 Similarity of the 32 plasmids envs and ONT sequencing statistics.**
(A) Accession numbers of the sequenced 32 envs. Note, ZM53M.PB12 has T1803G mutation in the plasmid compared to GenBank sequence. The mutation was confirmed by sanger sequencing.
(B) Heatmap to show sequence identity between the 32 plasmid envs.
(C) ENV sequencing run statistics using different basecalling model and read types.
(D) Raw read frequencies of the 32 references in four datasets. Blastn was used to assign each raw read to one of the 32 reference genes.

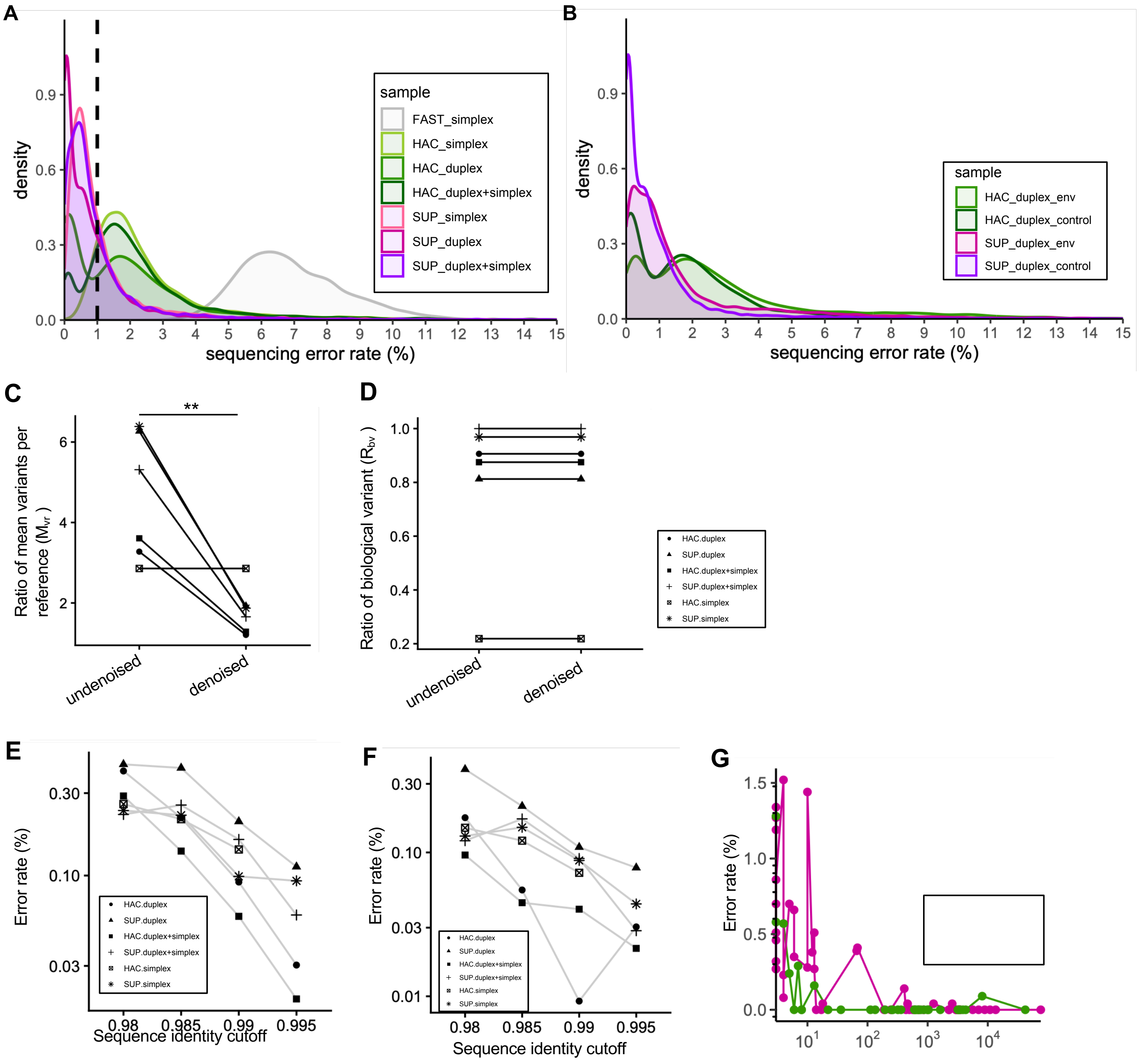


**Figure.S2 Sequencing quality and env variant calling of 32 strains.**

(A) Comparison of sequencing error rates of spikein lambda genome from duplex and simplex reads from fast, HAC, and SUP basecalling models. Reads with error rates higher than 15% were not shown.

(B) Comparison of sequencing errors of HIV-1 env and lambda genome. Reads with error rates higher than 15% were not shown.

(C) Denoise and chimera filter substantially reduced the ratio of mean variants per reference ($M_{vr}$). Usearch clustering with identity cutoff of 0.99 was used to generate consensus sequences. Paired T-test : p-value = 0.01.

(D) Denoise and chimera filter has no effect on the ratio of biological variants ($R_{bv}$). Usearch clustering with identity cutoff of 0.99 was used to generate consensus sequences.

(E) Error rates of curated envs from random subsampling of 100,000 duplex reads from HAC and SUP basecalled datasets. Error rates were calculated for curated envs with >=3, >=10, and >=15 read coverage. Ten subsampling repeats were generated.

(F) Error rates of curated envs from random subsampling of 150,000 duplex reads from HAC and SUP basecalled datasets. Error rates were calculated for curated envs with >=3, >=10, and >=15 read coverage. Ten subsampling repeats were generated.

(G) Sequencing error is reduced when sequencing depth increase. For both SUP and HAC reads, sequencing error is reduced substantially when sequencing depth increase to over 10 reads.

**, T-test p-value<0.01; ns, not significant.

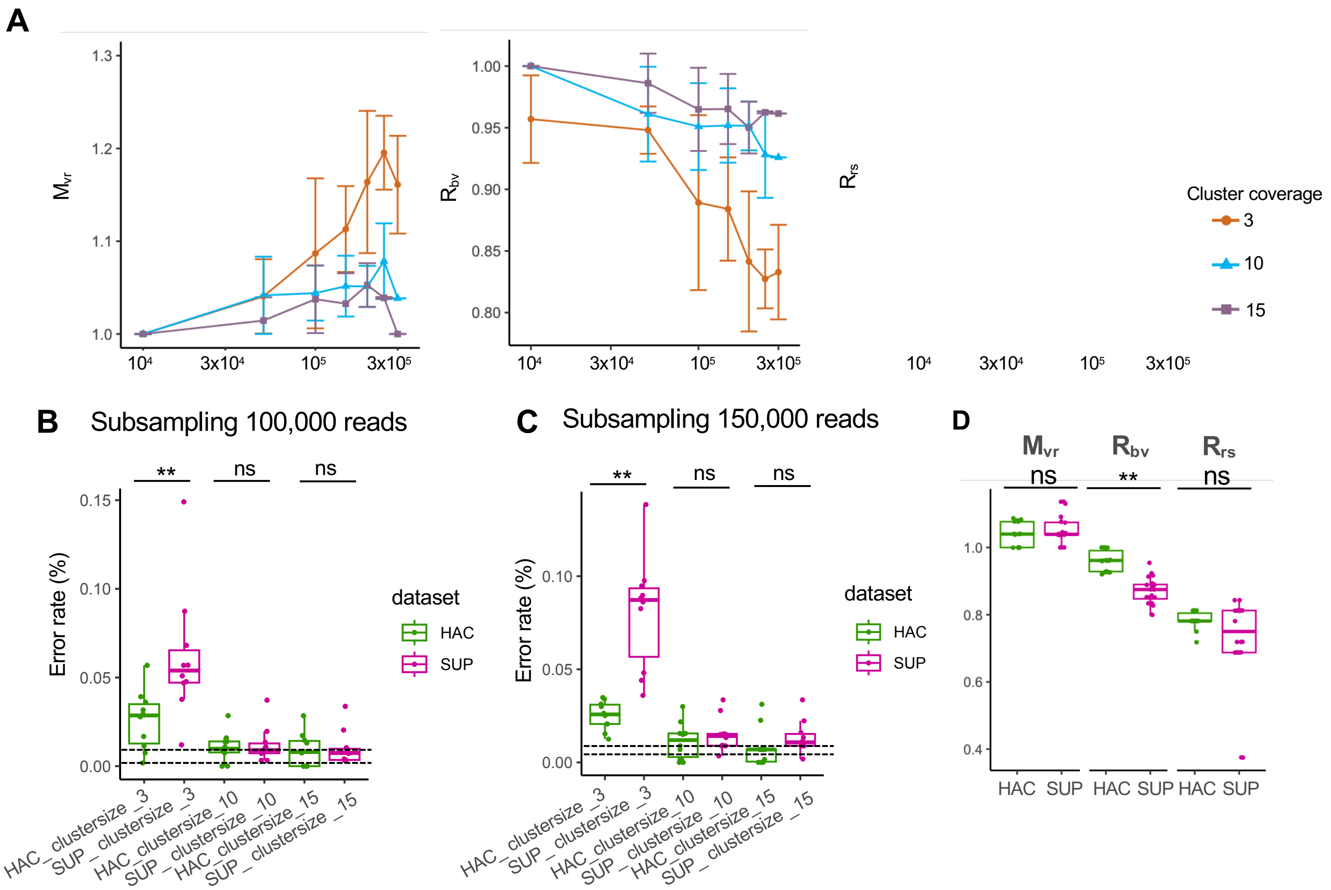


**Figure.S3 Sequencing quality and env variant calling of 32 strains.**

(A) Random subsampling revealed the distributions of $R_{bv}$, $R_{rs}$, and $M_{vr}$ with ONT duplex reads increasing from 10,000 to 300,000.

(B) Error rates of curated envs from random subsampling of 100,000 duplex reads from HAC and SUP basecalled datasets. Error rates were calculated for curated envs with >=3, >=10, and >=15 read coverage. Ten subsampling repeats were generated.

(C) Error rates of curated envs from random subsampling of 150,000 duplex reads from HAC and SUP basecalled datasets. Error rates were calculated for curated envs with >=3, >=10, and >=15 read coverage. Ten subsampling repeats were generated.

(D) Comparison of $R_{bv}$, $R_{rs}$, and $M_{vr}$ between HAC and SUP duplex datasets.

**, T-test p value<0.01; ns, not significant.

A

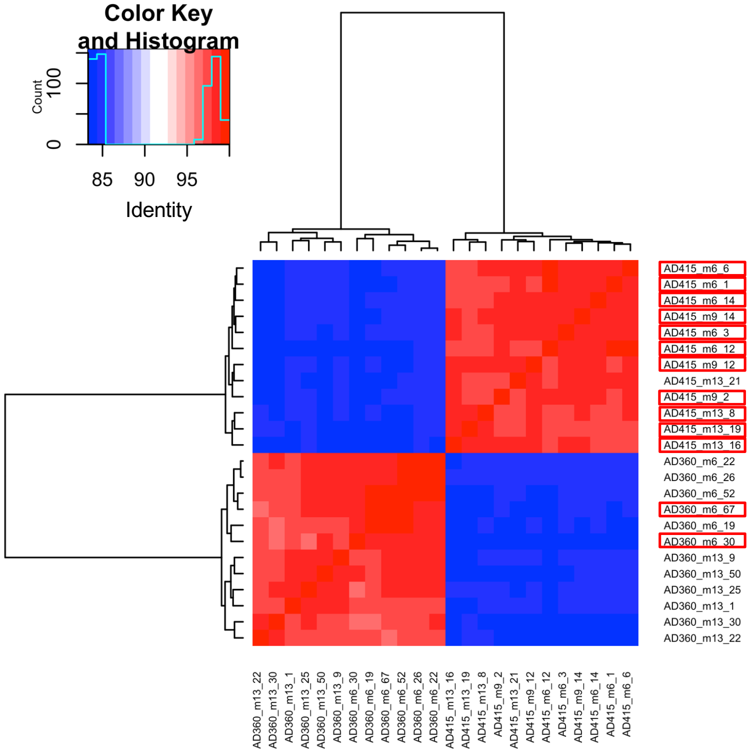


B

| sampleID | Raw reads | ENV_raw | ENV_unique |
| --- | --- | --- | --- |
| repeate1 | 115189 | 49015 | 26477 |
| repeate2 | 89846 | 38903 | 25963 |
| repeate3 | 117919 | 17397 | 10752 |

C

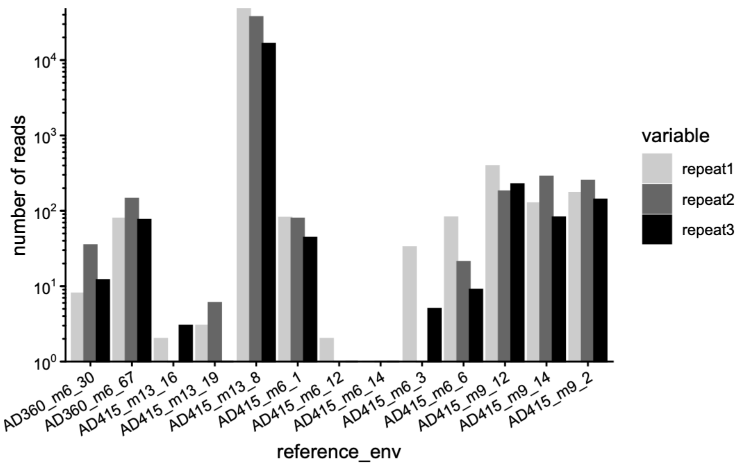


**Figure.S4 Sequencing quality and env variant calling of 30 SGA.**

(A) Sequence identity between env SGAs. SGAs detected in ONT datasets were highlighted in red.

(B) ENV sequencing run statistics. Unique reads are from usearch clustering with 0.99 identity. ONT reads are generated using duplex basecalling with the HAC model.

(C) Raw read frequencies of the 13 reference SGAs detected in three ONT repeats. Blastn was used to assign each raw reads to one of the 13 reference genes.

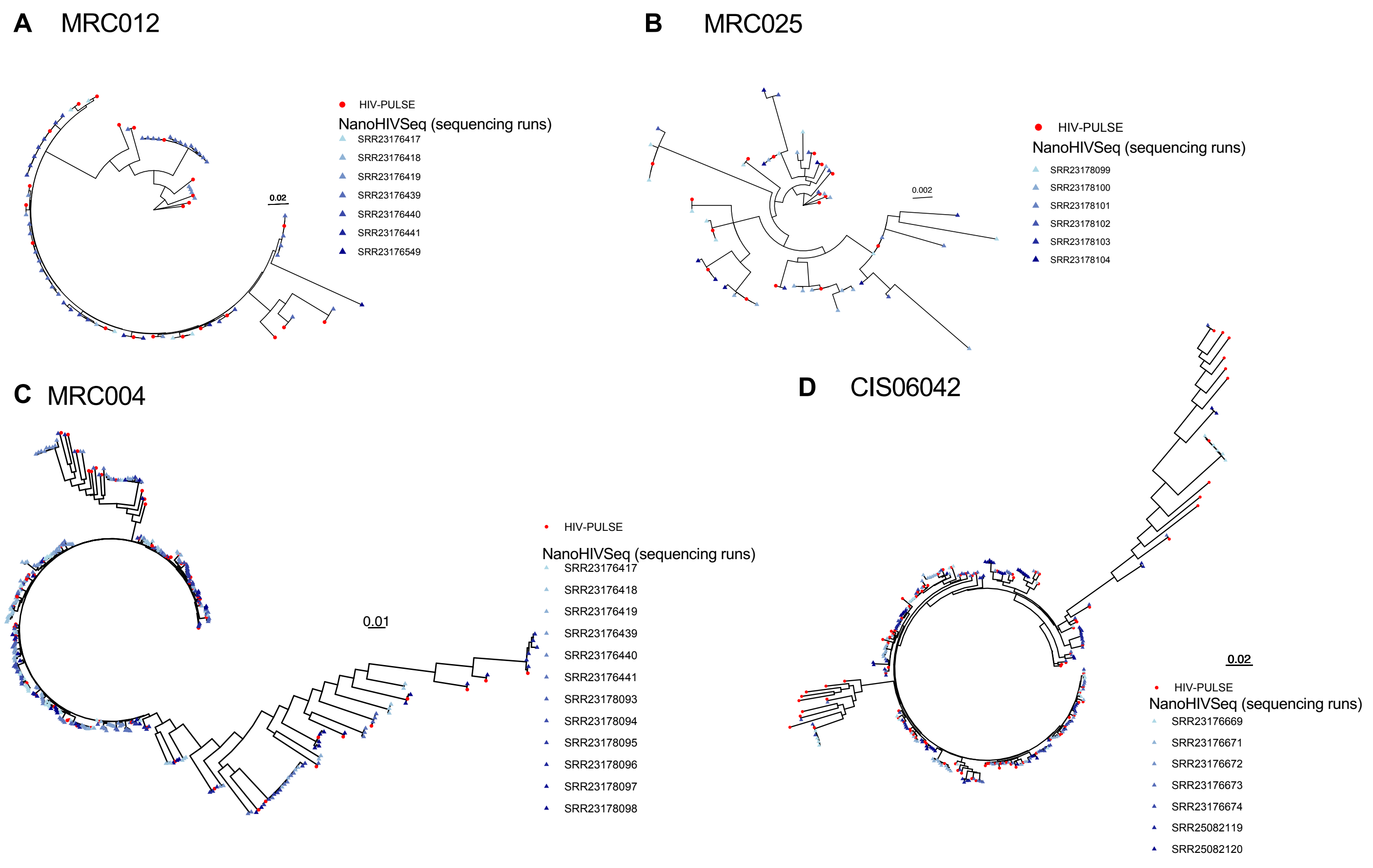


**Figure.S5 Comparison of HIV-PULSE and NanoHIVSeq-derived unique env sequences.**
(A) Donor MRC012 (B) Donor MRC025, (C) Donor MRC004, (D) Donor CIS06042. Each donor were sequenced multiple ONT runs whose SRA accession numbers are listed. NanoHIVSeq processed each ONT run separately. HIV-PULSE data are obtained from previous publication. The SRA run IDs were shown on the right.
